# Anti-diabetic effects of *Holarrhena antidysentrica* extracts: Results from a Longitudinal Meta-analysis

**DOI:** 10.1101/2021.02.11.430868

**Authors:** C. A. Divya, Sujan K Dhar, Manjula Shantaram, Manjula Das

**Author notes:** Equal Contribution (C. A. Divya, Sujan K Dhar). **E-mail addresses:** C. A. Divya, Sujan K Dhar, Manjula Shantaram, Manjula Das. **Corresponding Author:** Manjula Das, Mazumdar Shaw Medical Foundation, 8^th^ floor MSMC, Narayana Health City, Bommasandra, Bangalore 560099, India.

## Abstract

**Background:** *Holarrhena antidysenterica* (HA), a twining shrub belonging to the Apocynaceae family is found in tropical regions of Africa and over a large part of Asia including India, Philippines and Malayan Peninsula. In Indian traditional system of medicine, HA has been used to treat gastric ailments, for wound healing and also to improve glycaemic control. Glucose lowering activity of HA root, bark, seed, leaf and fruit extract in different parts of India as well as in Chinese traditional medicine is widely reported.

**Purpose:** In the meta-analysis reported in this article, we summarize glucose-lowering effects of HA extracts from different plant parts as reported in multiple studies involving animal models of diabetes. Our analysis helps to quantify the glucose-lowering effect of HA in comparison with standard diabetes drugs. The analysis also sheds light on differential efficacy levels of HA extracted from different plant parts.

**Study design:** The meta-analysis was carried out following PRISMA guidelines. Literature was searched to identify studies published between years 2011 to 2019 reporting glucose-lowering effects of HA extract on rodent models of diabetes.

**Methods:** Longitudinal meta-analysis was carried out on time-course data extracted from selected studies to calculate standardized mean change of glucose value from day 1 to days 7, 14 and 21 post-treatment by HA extract or standard anti-diabetic drug. Subgroup analysis was carried out for studies reporting effects of HA on leaf and seed extracts. Standardized mean difference in levels of cholesterol, triglycerides and serum total protein between treatment and control groups were also assessed.

**Results:** We shortlisted nine articles to be used for this meta-analysis. Summarized standardized mean changes of glucose value between day 1 and day 21 post-treatment indicated glucose-lowering effects of HA extracts to be marginally lower but comparable to that of standard anti-diabetic drugs like Glibenclamide or Sitagliptin. However, subgroup analysis revealed seed extracts of HA to be more potent than leaf extracts or even standard drugs. Effects of the extract on levels of cholesterol, triglyceride and serum total protein was also commensurate with its glucose-lowering property.

**Conclusions:** Our results, summarized over multiple studies, present a clear quantitative assessment of the anti-diabetic property of HA, in particular the seed extracts compared to standard anti-diabetic drugs. Further differential analysis of the seed extracts will be useful to arrive at a herbal formulation with superior anti-diabetic property and possibly lesser side effects than chemical entities.

## Introduction

Diabetes mellitus is a chronic and serious metabolic disorder primarily manifested with a hyperglycaemic condition as body is not able to produce sufficient quantity of Insulin hormone or is not able to utilize the produced hormone. Latest Diabetic Atlas of International Diabetic Foundation estimates around 463 million adults (20-79 years) living with diabetes worldwide which is an astounding proportion of 9.3% of global adult population(1). China, India and USA currently top the list of number of people with diabetes and are likely to retain their unenviable ranking in the next ten years. Mortalities in adults due to diabetes in 2019 was estimated as high as 4.2 million that is nearly twice the 2.11 million deaths recorded due to COVID-19 till date(2).

Several classes of non-insulin pharmacological agents including sulfonylureas, biguanides, thiazolidinediones, alpha-glucosidase inhibitors, dipeptidyl peptidase IV inhibitors are currently used in clinic to improve glycaemic control. However, undesirable side effects of many of these drugs often limit their use. On the other hand, natural product-based anti-diabetics especially medicinal plants and herbal formulations are very popular in many geographies due to their easy access, lesser cost and lesser side effects. Several medicinal plants from middle-east Asia, China, south-east Asia including India, Malaysia, Bangladesh, parts of Africa and Turkey have been documented in literature for their use as anti-diabetics(3). In particular India has a large repertoire of medicinal plants many of which have been used in traditional clinical practice to achieve glycaemic control(4,5). Herbal formulations from these medicinal plants(6) and phytocompounds derived from them (7) are shown to have glucose-lowering properties which make them interesting alternatives for chronic use.

*Holarrhena antidysenterica* (HA), an Indian medicinal twining shrub, belonging to the family Apocynaceae and commonly known as *Kurch* or *Kutaja* has been in use for long in folklore therapy(8). HA is a major ingredient in several Ayurvedic preparations such as *Bhunimbadi churna, Kutajghan Vati, Kutajarista* and *Kutaja churna,* that are used to treat dysentery, diarrhoea, fever and bacterial infections traditionally in India. In modern Ayurveda, HA is suggested for obesity, asthma, bronchopneumonia, hepatosplenomegaly, rheumatism(9) diabetes, cancer and even for wound healing(10). It is also used as an anti-oxidant and anti-depressant(11).

HA is studied extensively for treatment of metabolic diseases where the seed extract has shown alpha-glucosidase activity in *in vitro* assay(12). Ethanolic and methanolic extracts of bark from HA shown inhibitory effect of glycemia(13). Treatment of Streptozotocin (STZ) induced diabetic rats or mice by extracts of HA from different plant parts including seed, leaf and bark have resulted in lowering of blood glucose over a period of 7 to 21 days post-treatment(14–18)

Though many studies have indicated possible glucose-lowering and anti-diabetic activity of HA extract, its usefulness in comparison with known anti-diabetic compounds is not well established. Further, usage of various plant parts such as leaf and seed extracted with different solvent adds heterogeneity and ambiguity in results. In this article, we attempt to answer some of these questions by carrying out a careful meta-analysis of data collected from literature on observed anti-diabetic effects of HA extracts in animal models of diabetes in comparison with effects caused by standard anti-diabetic drugs like Glibenclamide or Sitagliptin. With a number of studies reported in literature to assess the anti-diabetic effects of HA extracts, to our knowledge this is the only meta-analysis study aiming to compare summary effects of HA extract to the effect caused by standard drugs.

## Meta-analysis Methods

### Selection of Literature

We performed the meta-analysis following PRISMA guidelines(19). Literature search was performed in PubMed and Google Scholar for articles in English published between 2011 and 2019 with search terms *Holarrhena antidysenterica* in title or abstract. Studies with STZ-induced diabetic rodent models were identified and included in meta-analysis. Review articles, commentaries, opinions, individual case studies and articles with only *in vitro* study results were excluded from analysis.

### Data Extraction

Selected articles were reviewed independently by two authors (CAD and SKD). Values of blood glucose and other parameters were extracted from articles using standardized forms and was cross-validated. Standard errors of various parameters reported in all selected studies except Keshri et al (20) were converted to standard deviation. Blood glucose values reported in mmol/L unit (16) were converted to mg/dL using the relationship 1 mmol/L = 18.018 mg/dL (21). Dose values of extracts and reference drug used, plant parts and solvents for extraction were recorded as mentioned in the articles. One article (22) did not provide the data in tabular form, for which, data was extracted from the graphs using WebPlotDigitizer online tool (23). For blood glucose time profile, the start time was normalized to the day of administration of extract or reference drug.

### Meta-Analysis and Meta-Regression

Choice of appropriate statistics for longitudinal meta-analysis is crucial. To measure the effect in change in blood glucose levels within the same group of animals we use a random effects model on the standardized mean change (SMC) with raw score standardization(24), between start of the treatment (day 1) and days 7, 14 and 21 post-treatment separately for animals treated with HA extract or reference drug. For other parameters, such as total cholesterol, triglycerides or serum protein measured at the end of the study period, we use the standardized mean difference (SMD) of measured parameter between treatment and diabetes control groups with Hedges’ correction for positive bias. All calculations were carried out using the Metafor library (25) on R statistical software platform. Meta-regression of SMC of blood glucose between start and day 21 of treatment was carried out using mixed-effects model with dose of the extract (in mg/kg) and other categorical variables such as plant part and solvent used for extraction as moderators.

## Results

### Study Details

The search yielded a total of 104 “hits” which were screened manually to select 9 articles containing levels of blood glucose and other parameters measured in animal models of diabetes (Fig. 1). All studies employed STZ-induced rodent models as the experimental platform.

**Figure 1:**
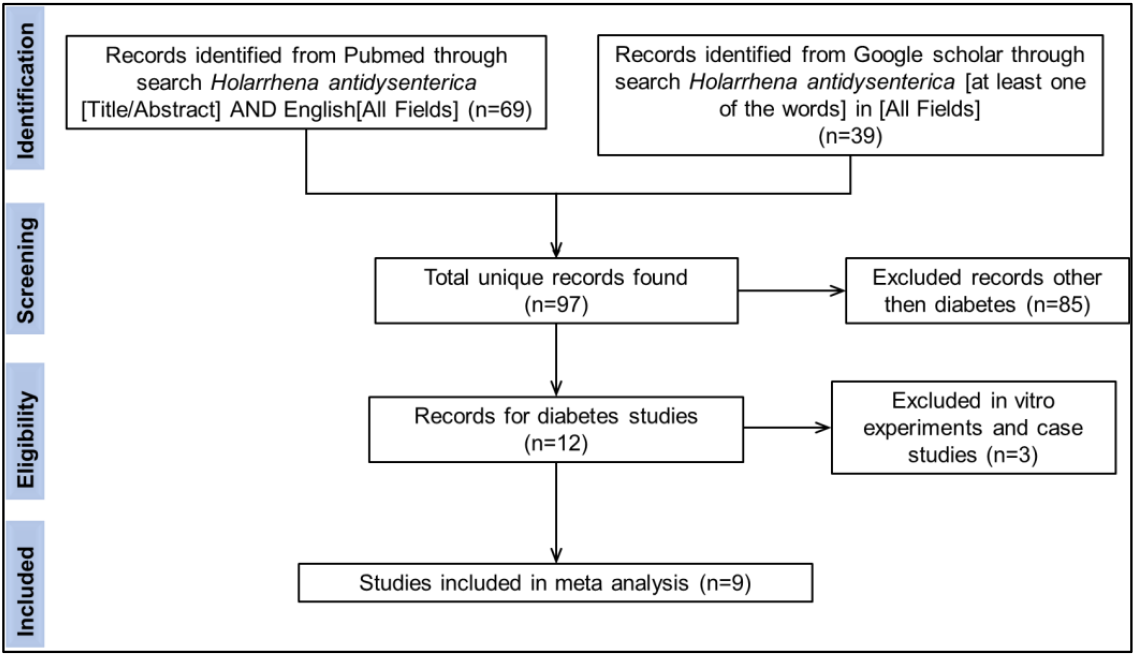
PRISMA flow diagram for literature search

Different plant parts of *H. antidysenterica* such as leaf (18,22) and seed (15,16,20,26–29) extracted with different solvents such as Ethanol, Ethyl acetate, Water, Petroleum ether and Methanol were used for treatment of the animals. Leaf or seed were powdered after air drying in shade and were extracted in a Soxhlet apparatus, employing respective solvents. Concentrated extract was air dried at room temperature or under reduced pressure and stored in air tight container in 2–8°C for use in experiments.

Some studies also compared glucose-lowering effects of standard drugs such as Glibenclamide (16,18,20,26–28) or Sitagliptin (22) along with HA on same experimental platform. Diabetes was induced by single intraperitoneal injection of STZ 50 mg/kg. Development of diabetes was confirmed by fasting blood glucose estimation 72 h post-STZ injection, wherein animals were fasted overnight before blood collection. Animals with fasting blood glucose level above 200 mg/dL at 72 h after STZ injection were considered diabetic and were included in experiments. Animals were randomly divided into groups of 6 each and assigned for treatment. Plant extract was administered orally every day in the dose range 100 – 600 mg/kg along with the reference drug at working concentration from the day of induction of diabetes. Fasting blood glucose was measured on days 1, 7, 14, 21 and 28 from start of the treatment.

Apart from serum glucose, most studies also reported values of other parameters (Table 1) out of which only three parameters – serum cholesterol, triglyceride and total protein were selected for meta-analysis since their values were reported in three or more studies.

**Table 1:** Baseline characteristics of selected studies

| Reference | Plant part | Solvent | Animal Model | No of Animals | HA dose (mg/kg) | Ref drug used | Ref drug dose (mg/kg) | Glucose measurement day | Other parameters measured |
| --- | --- | --- | --- | --- | --- | --- | --- | --- | --- |
| (18) | Leaf | Ethanol | STZ induced Wister albino rats | 6 | 200,400 | Glibenclamide | 5 | 1,7,14,21 | Body weight, GPT and GOT |
| (26) | Seed | Ethyl acetate | STZ induced Wister albino rats | 6 | Not mentioned | Glibenclamide | 0.6 | 1,7,14,21,28 | Triglyceride, HDL, Cholesterol, LDL, VLDL, Protein, Albumin, Insulin, body weight, Hepato-somatic index, Reno-somatic index, Globulin, Uric acid, Creatinine, Urea and Blood urea nitrogen. |
| (16) | Seed | 50% ethanol | STZ induced diabetic Sprague-Dawley rats | 6 | 100, 300 | Glibenclamide | 1 | 1,7,14,21 | Triglyceride, OGTT, Cholesterol, Lipids |
| (28) | Seed | Water, petroleum ether, methanol | STZ-induced diabetic Male wistar albino rats | 6 | 250 | Glibenclamide | 10 | Not measured | Triglyceride, Cholesterol, Protein, body weight, Urea |
| (29) | Seed | Water | STZ induced diabetic wistar strain male albino rats | 6 | 400 | Glibenclamide | 0.6 | 1,7,14,21,28 | Haemoglobin, Glycosylated Hb, Insulin, Body weight, creatinine, urea, GPT, GOT, ALP, |
| (22) | Leaf | Ethyl acetate | STZ-induced diabetic Wistar rats | 6 | 100, 200, 400 | Sitagliptin | 3 | 1,7,14,21,28 | Triglycerides, HDL, OGTT, Cholesterol, LDL, VLDL, Haemoglobin, Glycosylated Hb, Liver and Skeletal muscle glycogen, albumin, Insulin, Body weight, food and water intake, |
| (27) | Seed | 90 % Ethanol | STZ-induced diabetic Male albino wister rat | 6 | 300 | Glibenclamide | 5 | 1,7,14,21,28 | Body weight |
| (15) | Seed | Water | STZ-induced diabetic wister male albino rats | 6 | 600 | Not mentioned |  | 1,7,14 | Triglycerides, HDL, LDL, VLDL, Liver and Skeletal muscle glycogen, body weight, hexokinase, glucose-6-phosphatase, glucose-6-phosphate dehydrogenase, Liver and kidney GOT and GPT. |
| (20) | Seed | 90 % Ethanol | STZ-induced diabetic albino rats | 6 | 300, 600 | Glibenclamide | 5 | 1,7,14,21,28 | Triglycerides, OGTT, Protein, body weight, uric acid, creatinine, Urea, AST, ALP, ALK |

STZ: Streptozotocin, GPT: Glutamic Pyruvic Transaminase, GOT: Glutamic Oxaloacetic Transaminase, HDL: High Density lipoprotein, LDL: Low Density lipoprotein, VLDL: Very Low-Density lipoprotein, OGTT: Oral Glucose Tolerance Test, ALP/ALK: Alkaline phosphatase, AST: aspartate transaminase and ALT: alanine transaminase.

### Longitudinal Meta-analysis of Blood Glucose Levels

From results of the studies, it is seen that administration of HA extract has glucose-lowering effect over days 7 – 21 (Fig. 2). Results of longitudinal meta-analysis (Table 2) reveal that the effect estimated as SMC (difference with day 1) increases progressively from day 7 to day 21 for both HA extract as well as reference drug. On day 21, the effect size of reference drug (12.93, 95% CI: 6.44, 19.42) is higher than the effect produced by HA extract (9.53, 95% CI: 4.35, 14.70) (Fig. 3). However, results for HA extract showed higher degree of heterogeneity (I^2^ = 98.2%) compared to reference drug (87.3%) indicating possible variations in the effect size of HA extract due to dose, plant part or extraction solvent used.

**Figure 2:**
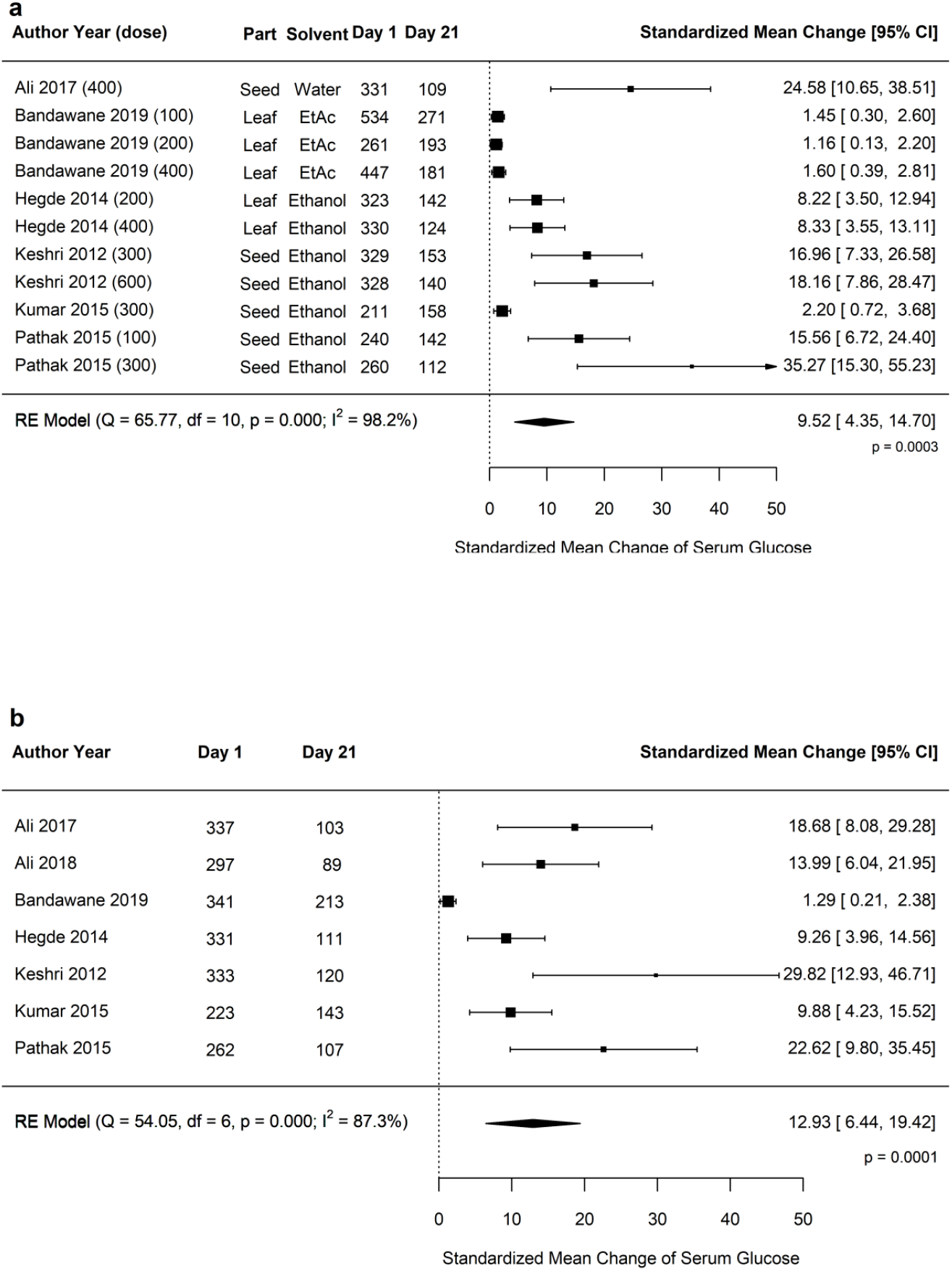
Effect on serum glucose by administration of HA extract

**Figure 3:**
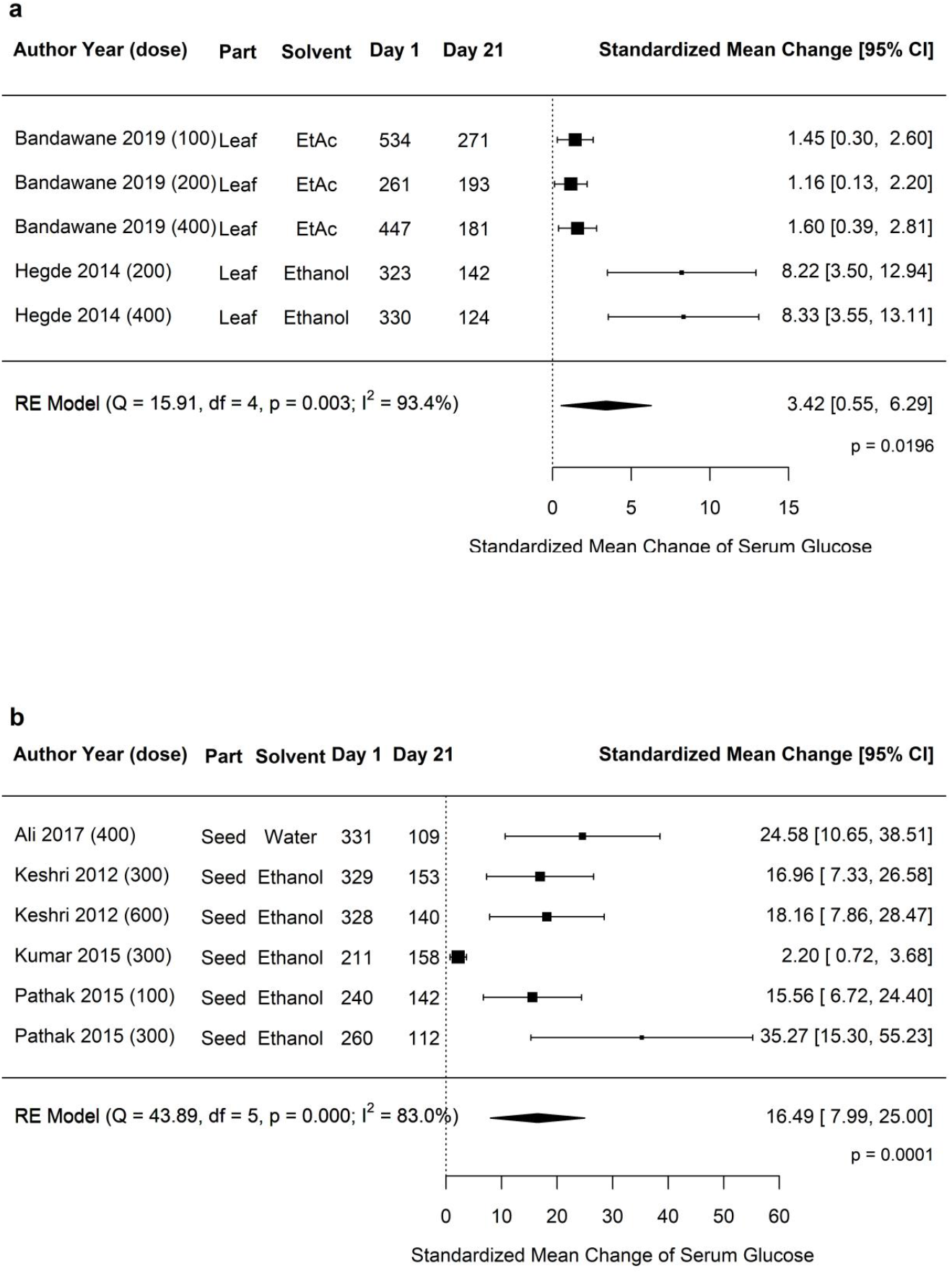
Standardized mean change of serum in animals treated with (a) HA extract and (b) reference drug glucose between day 1 and day 21 post-treatment

**Table 2:** Results of longitudinal meta-analysis of serum glucose after treatment with HA extract or reference drug

| Day after Treatment | Treatment with HA Extract |  | Treatment with reference drug |  |
| --- | --- | --- | --- | --- |
|  | Effect size (SMCR) on serum glucose with day 1 (95% CI) | Heterogeneity (I<sup>2</sup>) | Effect size (SMCR) on serum glucose with day 1 (95% CI) | Heterogeneity (I<sup>2</sup>) |
| Day 7 | 4.04 (1.41, 6.68)<br>p < 0.01 | 97.2% | 6.65 (2.97, 10.34)<br>p < 0.001 | 89.3% |
| Day 14 | 7.18 (3.02, 11.34)<br>p < 0.001 | 98.0% | 10.65 (4.95, 16.35)<br>p < 0.001 | 89.4% |
| Day 21 | 9.53 (4.35, 14.70)<br>p < 0.001 | 98.2% | 12.93 (6.44, 19.42)<br>p < 0.0001 | 87.3% |

### Meta-Regression and Subgroup Analysis

Meta-regression of the serum glucose SMC on day 21 for HA extract with dose as moderator did not show any statistically significant dependence of the SMC with dose of extract administered (Supplementary Fig. 1). To estimate variations due to plant parts, we performed the meta-analysis for the two subgroups, HA leaf and seed extracts used in studies. This subgroup analysis results (Fig. 4) indicate that extract from seeds have enhanced glucose-lowering ability (SMC 16.49, 95% CI: 7.99, 25.90) compared to that of leaf (SMC 3.42, 95% CI: 0.55, 6.29). It explains major portion of the observed heterogeneity of SMC of HA extract in Table 2. In all selected studies seeds were extracted in ethanol, whereas ethanol, ethyl acetate and water were used for extraction of leaves. Hence, we did not perform a separate subgroup analysis on the solvent as the results will not be different than results of plant part subgroups. Nevertheless, it is important to note that the seed extracts appear to be the most potent anti-diabetic fraction with a higher glucose lowering effect than the observed effects of Glibenclamide or Sitagliptin.

### Meta-analysis of other Markers

Meta-analysis of the other markers shows both HA extract and reference drugs reduce total cholesterol (Supplementary Fig. 2) and triglycerides (Supplementary Fig. 3) in serum on day 21 compared to the corresponding value in control diabetic animals who did not receive any treatment. SMD value for reference drugs is higher than that of HA extract (Table 3). However, for serum total protein, the reference drugs do not show any statistically significant summary effect (p > 0.1) whereas with HA extract a marginally significant effect (p = 0.051) is observed (Supplementary Fig. 4).

**Table 3:**
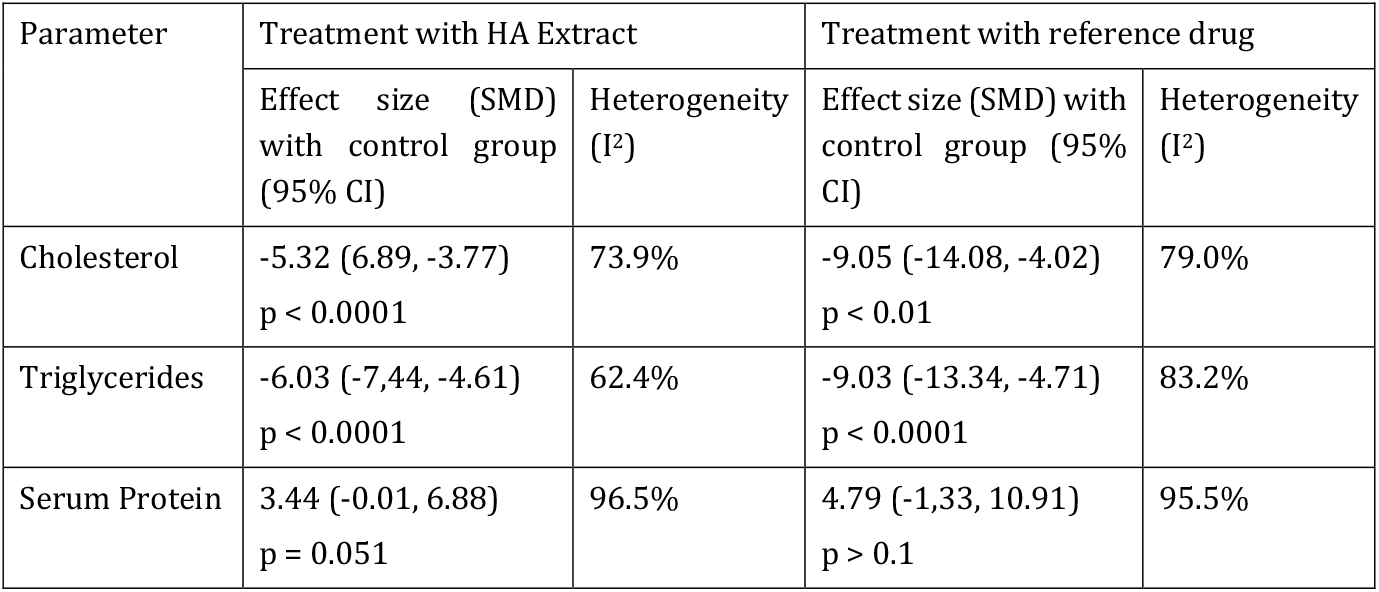
Meta-analysis results for the change of cholesterol, triglycerides and serum total protein between treatment and control group of animals

### Limitations

Primary limitation of this meta-analysis is the nonuniformity of plant parts and extraction solvent used across different studies. We performed subgroup analysis on the plant parts to assess their differential effects on glucose-lowering. However, similar subgroup analysis for extraction solvents could not be performed as ethanol was mostly used for extraction of seeds. Most studies used Glibenclamide as the reference drug except one, which compared the effect of Sitagliptin(22). However, for the comparison of anti-diabetic effect with HA extract, we summarized effects of both drugs together. All selected studies used the same number of animals (n = 6) for each treatment group, hence test for publication bias due to differing number of samples was not carried out.

## Discussions

Anti-diabetic properties of HA extract are well-known and is reported in many studies including the ones synthesized in this meta-analysis. However, the consolidation of data across multiple diverse studies clearly and quantitatively establishes the HA extract as a potent anti-diabetic agent comparable to standard drugs like Glibenclamide and Sitagliptin. Meta-analysis results for the effects of HA extract to lower the levels of serum cholesterol and triglyceride and restore levels of serum total protein towards normalcy further supports it’s efficacy as an anti-diabetic.

An important finding from this meta-analysis is the dose independence of HA extract to its observed glucose-lowering effect, that was not apparent from results of individual studies. However, this is not an uncommon observation, as dose independence of pharmacological effects of plant extracts were reported in studies with Musa AAA fruit (30), leaf extracts of *K. Africana(31)* and *Camellia sinensis* green tea extract(32). Possible explanation for this dose independent behaviour of HA extract could be the saturation of active transport of phytochemicals to cells in 100 – 600 mg/kg dose range with the excess amount getting excreted(33). Studies with HA extract at a lower dose range is recommended to identify its dose-dependent anti-diabetic activity.

It’s also important to note that seeds of HA extracted with ethanol showing markedly higher glucose-lowering effect. A comparison of available data on phytochemicals present in ethanol extract of seed and ethanol /ethyl acetate extracts of leaf (Table 4) shows differential presence of triterpinoids like betulinic acid, oleanolic acid and amyrin that are known anti-diabetic agents. STZ-induced diabetic mice treated with ß-amyrin showed reduction in the STZ-induced levels of blood glucose, cholesterol and triglycerides(34). Similar glucose lowering effect on a STZ-nicotinamide induced diabetic mice model was observed for treatment with betulinic acid(35). Oleanolic acid, another triterpenoid, was also shown to reduce blood glucose in STZ-induced rats. This anti-diabetic activity of oleanolic acid was associated with restoration of mRNA levels of anti-oxidant enzymes glutathione peroxidase 1 and superoxide dismutase 1 in liver of animals(36). It’s likely that the anti-oxidant properties of these triterpenoid molecules render their anti-diabetic potential. Quercetin, a flavonoid present in the seed extract is also studied extensively for its anti-diabetic potential. A recent meta-analysis shows glucose-lowering effect of quercetin in STZ or alloxan-treated diabetes model synthesized over 13 different studies(37).

**Table 4:** Comparison of phytochemicals present in ethanol extract of HA seed and ethanol / ethyl acetate extract of leaf

| Plant part | Solvent | Compound | References |
| --- | --- | --- | --- |
| Seed | 50 % Ethanol | alkaloids, carbohydrates, flavonoids including quercetin, tannins and phenolic compounds | (16) |
| Seed | 90 % Ethanol | Alkaloids, carbohydrates, flavonoids, tannins and phenolic compounds, Saponins and steroids | (27) |
| Seed | Ethanol | Serine protease, pentacyclic triterpenoids, (lupeol, betulinaldehyde, and betulinic acid) steroidal compound (stigma sterol), dihydrocanaric acid, amyrin, betulin, and oleanolic acid | (38) |
| Leaf | Ethanol | Flavonoids, alkaloids, tannins and steroids | (18) |
| Leaf | Ethanol | Saponins, Amino acids, Phenol and Glycosides | (39) |
| Leaf | Ethanol | Terpenoids and reducing sugars | (40) |
| Leaf | Ethyl acetate | Total phenols, tannin and flavonoids | (22) |
| Leaf | Ethyl acetate | Flavonoids Alkaloids, Glycosides, Terpenoids, reducing sugars, Saponins, Tannins and Steroids | (40) |

These results surely indicate the possibility of individual triterpinoids or flavonoids or their combinations to render the enhanced anti-diabetic potential of HA seed extracts. More detailed studies with differential identification of compounds between seed and leaf extracts will perhaps unlock the true antidiabetic potential of HA as a herbal formulation.

## Conclusion

This meta-analysis conclusively summarizes the anti-diabetic effect of HA extract seen in rodent diabetic models across 9 different studies. Ethanol extracts of HA seeds appear to have more potent glucose-lowering ability than reference drugs like Glibenclamide and Sitagliptin. Further, the HA extracts also lowered serum cholesterol and triglycerides and restored total serum proteins in diabetic animals. Further studies on HA extracts can lead to novel herbal formulations with anti-diabetic effects comparable to chemical compounds.

## Author contribution

CAD: data curation and manuscript preparation, SKD: data curation, statistical analyses and manuscript preparation, MD: conceived the project and manuscript review, MS: manuscript review.

## Acknowledgement

SKD acknowledges Prof. Joy Kuri, Chair, Department of Electronic Science and Engineering, Indian Institute of Science, Bangalore for providing the computational resources.

## Supplementary Material

Supplementary information file enclosed

## Abbreviations

CI: confidence interval
HA: Holarrhena antidysenterica
mRNA: messenger ribonucleic acid
PRISMA: preferred reporting items for systematic reviews and meta-analyses
SMC: standardized mean change
SMD: standardized mean difference
STZ: Streptozotocin

---

***Figure 4: Subgroup analysis of standardized mean change of serum glucose between day 1 and day 21 in animals treated with (a) HA leaf extract and (b) HA seed extract.***

