## Supplementary Information for "Anti-diabetic effects of *Holarrhena antidysentrica* extracts: Results from a Longitudinal Meta-analysis"

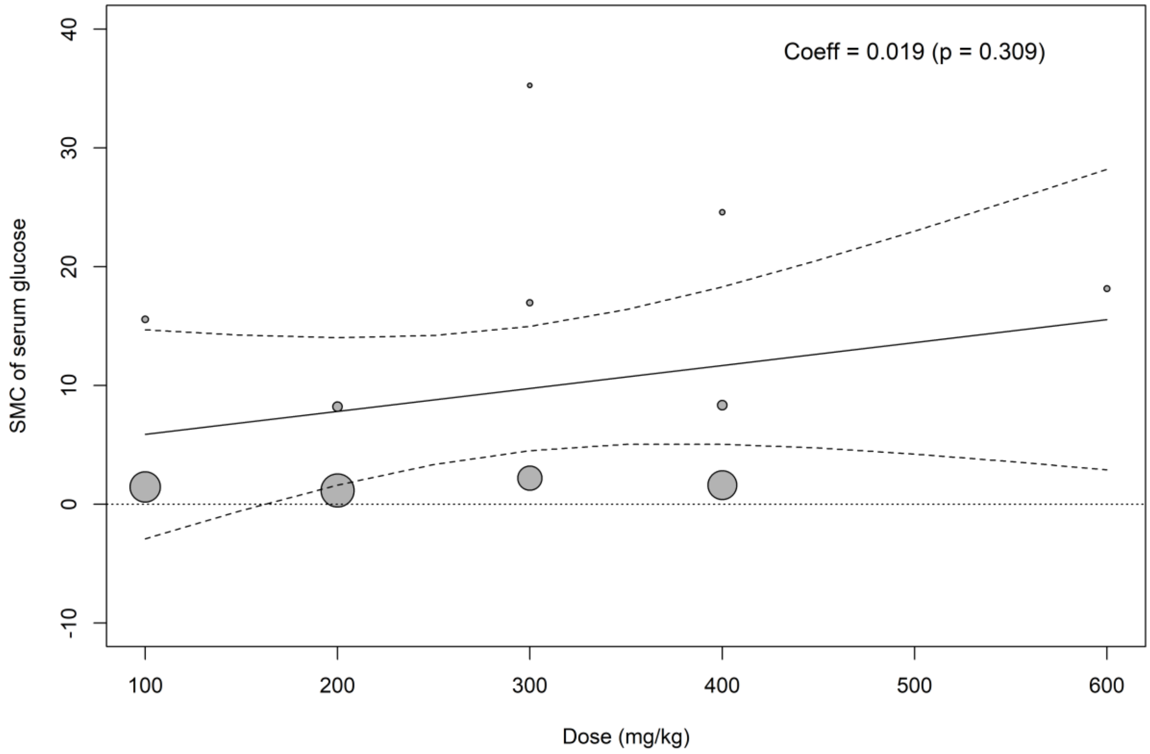


***Supplementary Figure 1: Meta-regression of standardized mean change of serum glucose between day 1 to day 21 post-treatment with dose of HA extract***

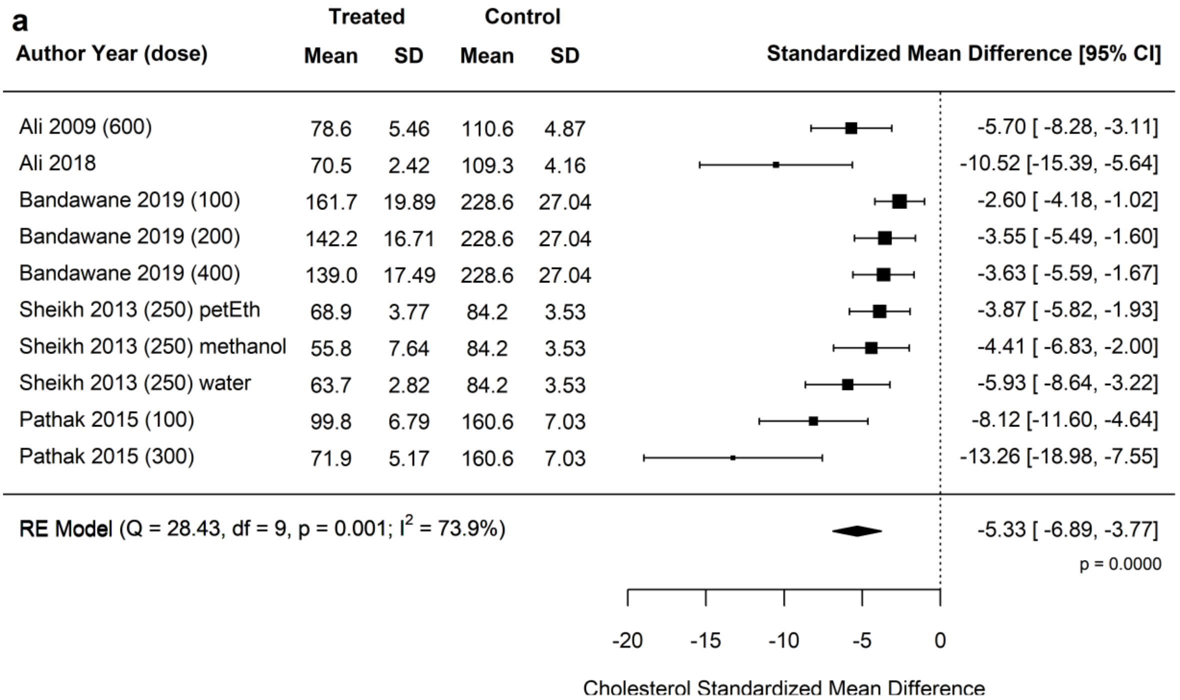


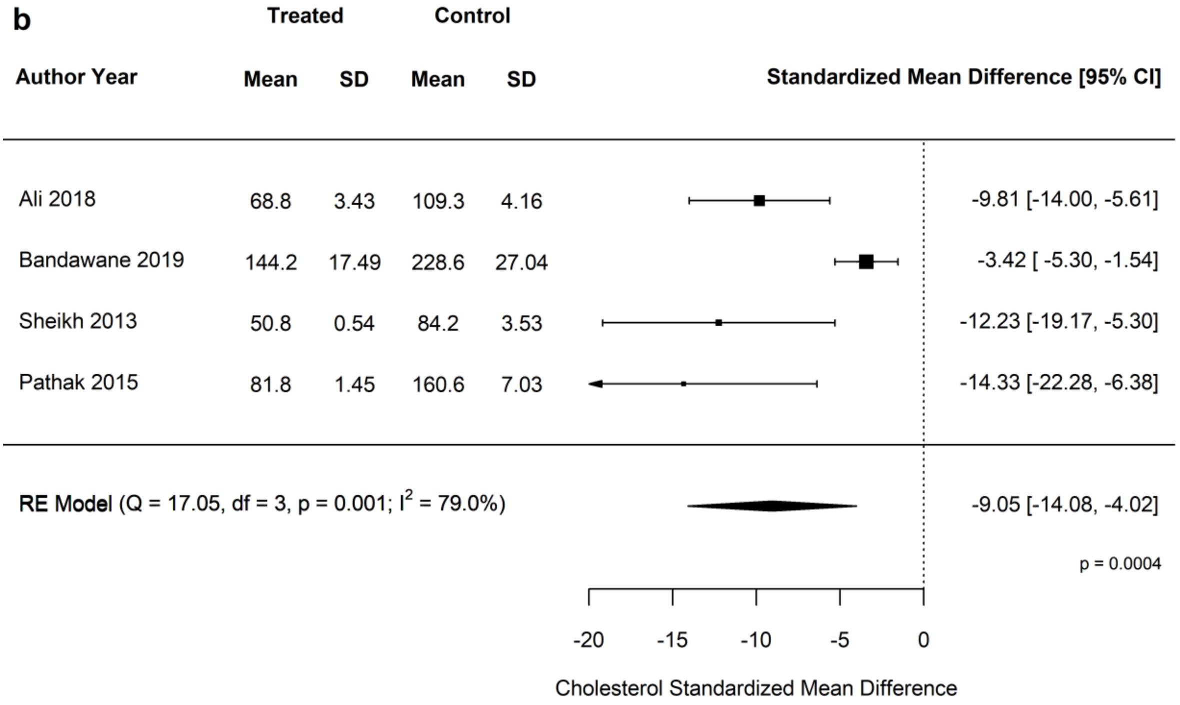


***Supplementary Figure 2: Standardized mean difference of serum cholesterol between diabetic animals treated with (a) HA extract and (b) reference drug and control group of animals***

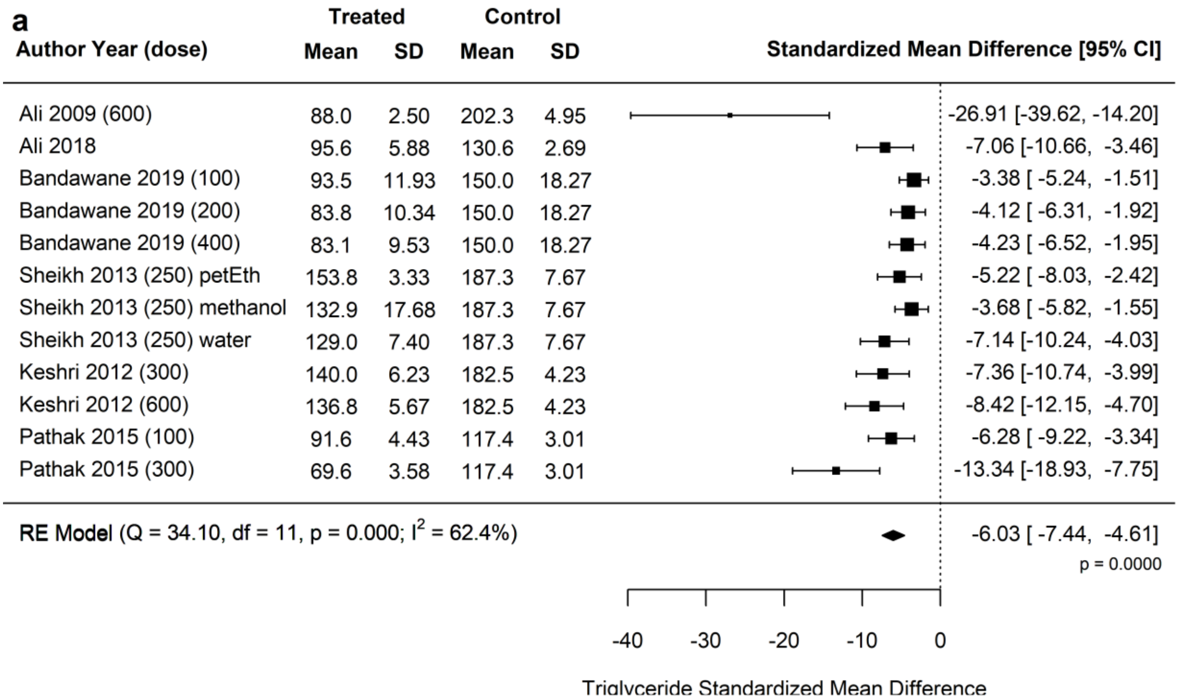


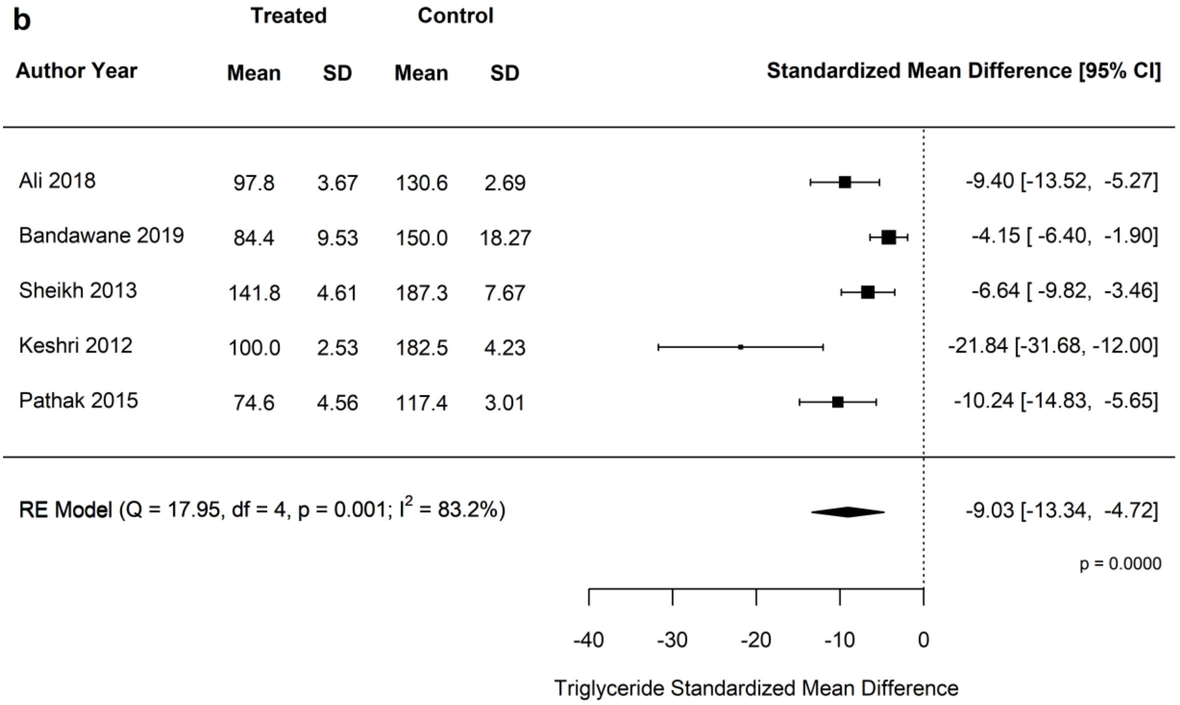


***Supplementary Figure 3: Standardized mean difference of serum triglycerides between diabetic animals treated with (a) HA extract and (b) reference drug and control group of animals***

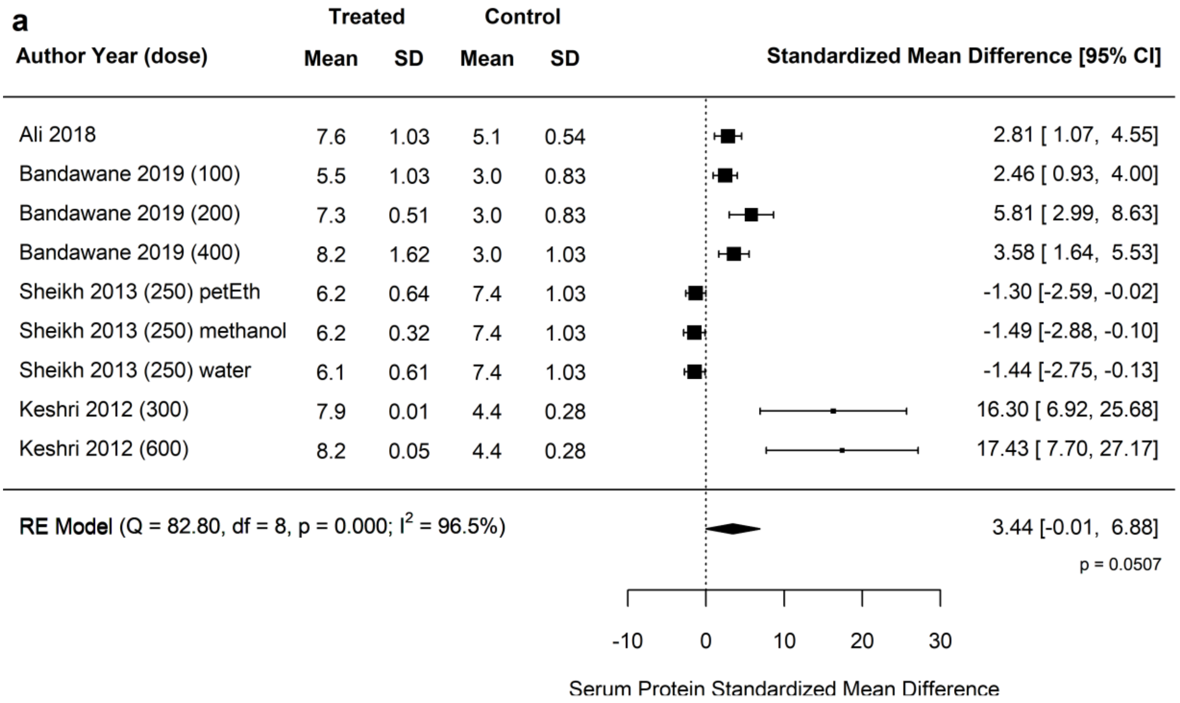


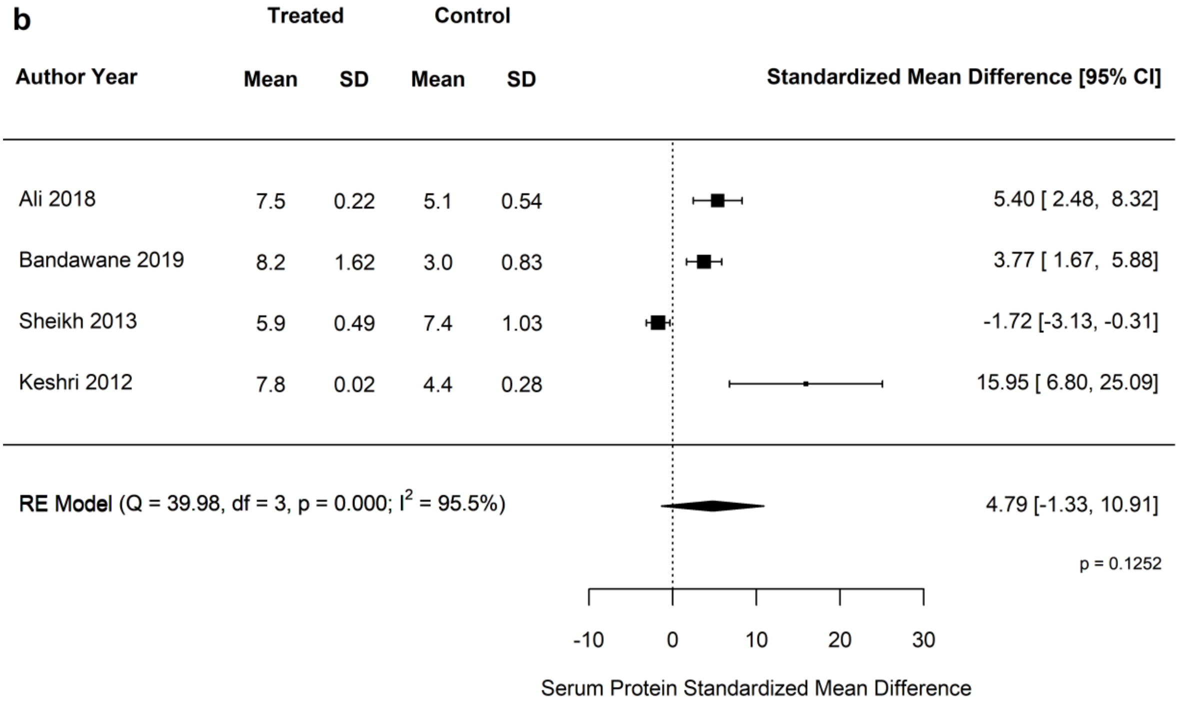


***Supplementary Figure 4: Standardized mean difference of serum total protein between diabetic animals treated with (a) HA extract and (b) reference drug and control group of animals***
